# Asymmetrically Positioned Flagellar Control Units Regulate Human Sperm Rotation

**DOI:** 10.1101/290122

**Authors:** Melissa R. Miller, Samuel J Kenny, Nadja Mannowetz, Steven A. Mansell, Michal Wojcik, Sarah Mendoza, Robert S. Zucker, Ke Xu, Polina V. Lishko

## Abstract

The ability of sperm to fertilize an egg is controlled by ion channels, one of which is the pH-dependent calcium channel of sperm CatSper. For CatSper to be fully activated, the cytoplasmic pH must be alkaline, which is accomplished by either proton transporters, or a faster mechanism, such as the voltage-gated proton channel Hv1. To ensure effective regulation, these channels and regulatory proteins must be tightly compartmentalized. Here, we characterize human sperm nanodomains that are comprised of Hv1, CatSper and regulatory protein ABHD2. Super-resolution microscopy revealed that Hv1 forms asymmetrically positioned bilaterally distributed longitudinal lines that span the entire length of the sperm tail. Such a distribution provides a direct structural basis for the selective activation of CatSper, and subsequent flagellar rotation along the long axis that, together with hyperactivated motility, enhances sperm fertility. Indeed, Hv1 inhibition leads to a decrease in sperm rotation. Thus, sperm ion channels are organized in distinct regulatory nanodomains that control hyperactivated motility and rotation.

## Introduction

Fast cellular responses, such as phototransduction or ciliary motility, rely on efficient signal transduction. To achieve rapid signaling, the members of the transduction cascade are tightly compartmentalized into regulatory nanodomains and are located in close proximity to each other (Burns and Pugh, 2010). The organization of the sperm flagellum should follow the same logic, as motility cues must propagate rapidly to achieve concerted movement. As mammalian spermatozoa ascend in fallopian tubes, they have to overcome upstream fluid flow. To succeed in such a task, spermatozoa engage in rheotaxis (Ishijima et al., 1992; Kantsler et al., 2014; Miki and Clapham, 2013), and display specific flagellar motility patterns known as hyperactivation and rolling, the latter characterized by a rotation of the flagellum around its longitudinal axis(Ishijima et al., 1992). Such rotation is thought to be linked to a calcium influx, and is required for successful rheotaxis (Miki and Clapham, 2013). Flagellar movement is generated by microtubule sliding powered by ATP hydrolysis (Lindemann and Gibbons, 1975) – a highly pH-dependent process, which could also be augmented by an elevation of intraflagellar calcium (Brokaw, 1979; Ho et al., 2002; Ishijima et al., 2006; Lindemann et al., 1987; Suarez, 2008; Suarez et al., 1993). Therefore, sperm intracellular calcium concentration [Ca^2+^]_i_ and pH, in addition to ATP, are the main regulatory mechanisms allowing for motility changes and are under control of ion channels and transporters (Lishko et al., 2012; Ren and Xia, 2010). Distinct nanoscale spatial organization of proton and calcium channels, then, may be a requirement for efficient sperm motion. The principal calcium channel of mammalian sperm CatSper (Carlson et al., 2005; Carlson et al., 2003; Chung et al., 2014; Kirichok et al., 2006; Liu et al., 2007; Qi et al., 2007; Quill et al., 2001; Ren et al., 2001; Smith et al., 2013; Xia et al., 2007) is regulated by intracellular alkalinization (Kirichok et al., 2006; Lishko et al., 2012), which, in turn, is under control of yet-to-be-identified proton exchangers and likely the voltage-gated proton channel Hv1 (Lishko et al., 2010). The latter mechanism is only reported for human spermatozoa, and is not detected in murine sperm (Lishko and Kirichok, 2010). Interestingly, while both human and mouse spermatozoa hyperactivate, only human spermatozoa display full 360-degree rotation along the long axis, while mouse sperm alternate between 180-degree turns(Babcock et al., 2014). CatSper, which is organized in quadrilateral longitudinal nanodomains along the sperm flagellum (Chung et al., 2017; Chung et al., 2014) is also regulated by species-specific cues (Miller et al., 2015). In human spermatozoa, CatSper is activated by both flagellar alkalinity and progesterone (Lishko et al., 2011), while in murine sperm, CatSper is activated solely by flagellar alkalinity (Kirichok et al., 2006). One candidate for proton extrusion in human sperm is the voltage-gated proton channel Hv1, which moves protons unidirectionally toward the extracellular space and raises intracellular pH (Ramsey et al., 2006; Sasaki et al., 2006). Hv1 is expressed in human spermatozoa (Lishko et al., 2010), where it can trigger intracellular alkalinization, ensuring a favorable condition for activation of the pH-sensitive calcium channel CatSper (Kirichok et al., 2006; Ren et al., 2001). Resulting calcium influx via CatSper directly affects axonemal function, and triggers change in the motility behavior known as hyperactivation and rolling, or rotation along the long axis. While both are needed for sperm fertility and to detach from the oviductal epithelia (Avenarius et al., 2009; Babcock et al., 2014; Carlson et al., 2005; Ho et al., 2002; Miki and Clapham, 2013), their precise regulating mechanisms have not been identified. Here, we investigated whether the specific localization and regulation of the sperm proton channel Hv1, as well as CatSper and its regulatory protein ABHD2 (Miller et al., 2016) could explain these specific motility patterns.

## Results

### Hv1 forms bilateral longitudinal lines that span the entire length of the sperm tail

Human sperm flagellum is structurally divided into three main regions: a short, mitochondria-rich midpiece, a longer principal piece, and a very short endpiece with few structural elements (Figure 1, bottom panel). According to our previously published Hv1 immunostaining in human spermatozoa (Lishko et al., 2010), Hv1 distribution is restricted to the principal piece of the flagellum, and this channel is functionally active in both epididymal and ejaculated human spermatozoa (Figure S1). However, traditional epi-fluorescence microscopy is not suitable for revealing the details of Hv1 pattern distribution due to its diffraction-limited resolution of ∼250 nm, while sperm flagellar diameters are on sub-micron scales. To overcome this, the three-dimensional stochastic optical reconstruction microscopy (3D STORM) (Huang et al., 2008; Rust et al., 2006) was used to reveal detailed Hv1 flagellar organization in human spermatozoa. Remarkably, 3D STORM images showed that Hv1 forms bilateral lines along the principal piece of the flagellum (Fig. 1 and Fig. S2), and such localization was preserved upon sperm in vitro capacitation (Fig. S3A). During such treatment Hv1 channel undergoes N-terminal cleavage (Berger et al., 2017); however, its distinct bilateral distribution pattern does not change (Fig. S3B*-C*). Interestingly, Hv1 bilateral distribution differs from the longitudinal quadrilateral arrangement of the murine and human calcium channel CatSper (Fig. S3*D*, and (Chung et al., 2017; Chung et al., 2014)). Detailed analysis of the distance between Hv1 bilateral lines gave an average distance of 228 ± 19 nm midway down the length of the sperm tail, with 230 ± 15 nm and 250 ± 12 nm for the endpiece region and for the proximal part of the principal piece, respectively (n=6 for each measurement; Fig. 2*A*). Interestingly, the flagellar diameter of human sperm is at least double the distance measured between the discrete Hv1 lines. The diameter of the human sperm flagellum varies in the literature from 400 to 600 nm; such a wide range of reported values is likely due to differences in imaging method and in electron microscopy sample preparation methods, which often distort the sample. A precise measurement of sperm diameter using particular imaging routine and pool of human donors used in this study was essential to the interpretation of this peculiar Hv1 distribution. Therefore, the diameter at the level of the principal piece was measured using brightfield, fluroescence, and electron microscopy techniques and found to be between 436 nm and 634 nm. Scanning electron microscopy (SEM) and differential interference contrast (DIC) images (Fig. S4*A-D*) indicated an average diameter of human sperm flagellum used in this study of 570±13 nm (n=3) and 555±10 nm (n=7), respectively. To corroborate this measurement, the sperm membrane protein ABHD2 (Miller et al., 2016) was immunolabeled for STORM imaging. Distance measurements taken from STORM images of ABHD2 at the level of the principal piece of human sperm tails returned similar values of: 452±37 nm, 431±13 nm, and 404±27 nm (corresponding to cross-sections of the proximal, middle, and distal positions as mentioned above for Hv1 (n=5); Figs. 1*C-D*, 3*C-D* and Fig. S4*E*).

**Figure 1.**
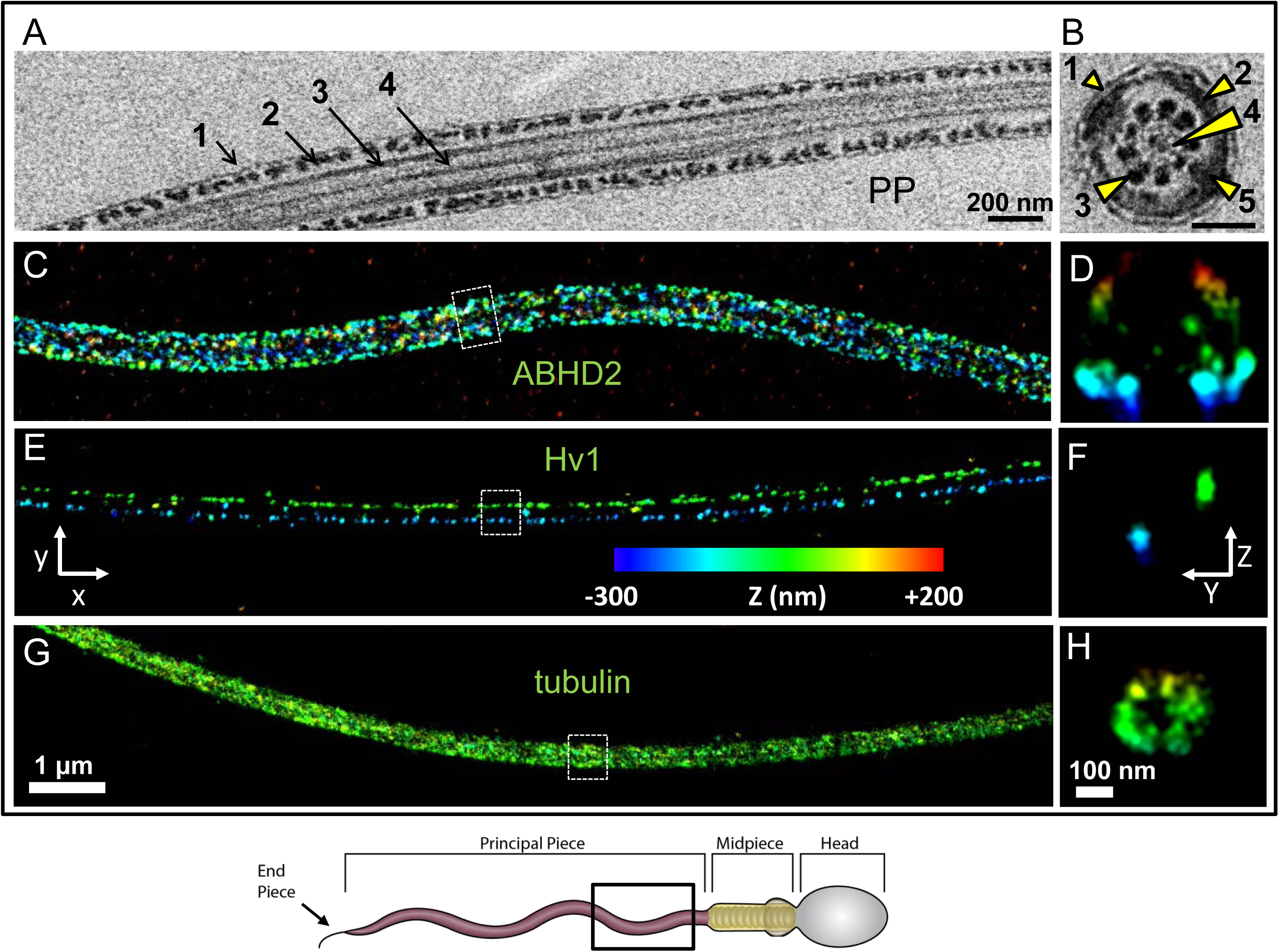
STORM reveals distinct localization of regulatory elements within human sperm cells. *Bottom panel:* schematic representation of spermatozoon with cellular compartments labeled. (A), (C), (E), and (G) panels show conventional x/y projections of the sperm flagellum, while (B), (D), (F) and (H) show cross-sections in y/z projections from the corresponding cells, except for panel (B), whose cross-section is from a different cell. (A) TEM of a human sperm principal piece (PP) with structural elements as: 1-plasma membrane; 2-fibrous sheath; 3-outer dense fibers; 4-microtubules. (B) Cross section of the flagellum at the level of PP. Elements are as in (a), as well as: 5-symmetrically positioned longitudinal columns. (C) Immunostaining for CatSper regulatory protein ABHD2. The corresponding cross section (D) of the boxed region reveals quadrilateral arrangement of ABHD2. (E) Immunostaining for Hv1 and the cross section (F) of the boxed region reveals that Hv1 forms bilateral lines. (G) Immunostaining for beta-tubulin and the cross section (H) of the boxed region used as a marker of the axoneme. Scale bars are 200 nm (A) and (B), 1 μm (C, E, and G) and 100 nm (D, F and H). The color in all projections reflects the relative distance from the focal plane along the z axis as shown on the color scale bar in (E).

**Figure 2.**
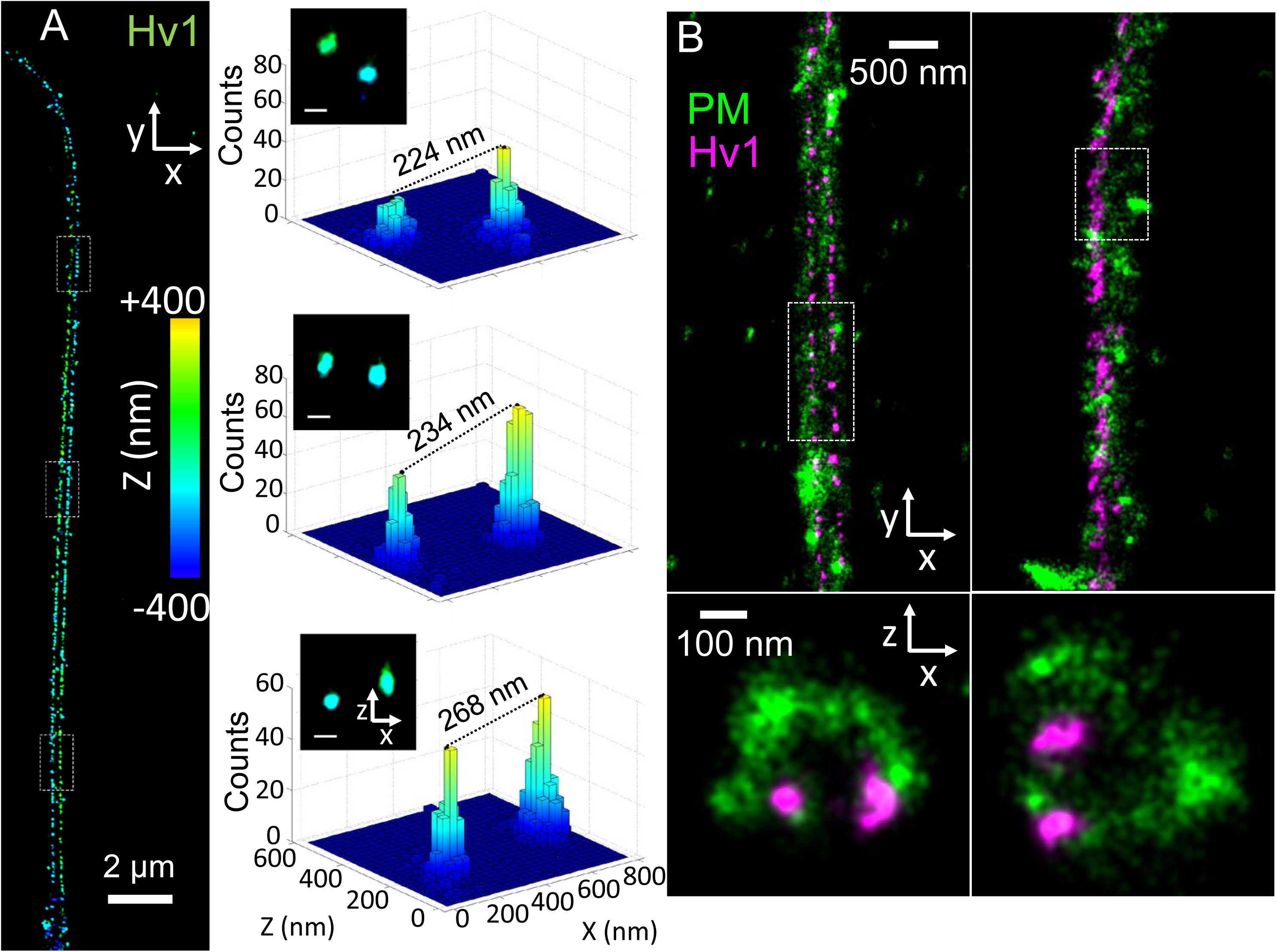
Human sperm Hv1 ion channels are organized into bilateral domains. (A) Left panel: super-resolution image of Hv1 in a representative human sperm flagellum. Right panel: three regions of interest (ROIs) from x/y projections (dotted boxes in the left panel) were examined with x/z projections shown. For each ROI, the distance between two Hv1 signals was measured as the peak to peak distance from 2D histograms of STORM localizations. (B) Two-color image of Hv1 (magenta) and the plasma membrane (green, stained with CM-DiI) in x/y and x/z projections. The x/z projections correspond to the boxes in the x/y projections. Shown are two representative flagella. Scale bars: 2 μm ((A), left), 100 nm ((A), right inset; (B), lower), and 500 nm ((B), upper).

**Figure 3.**
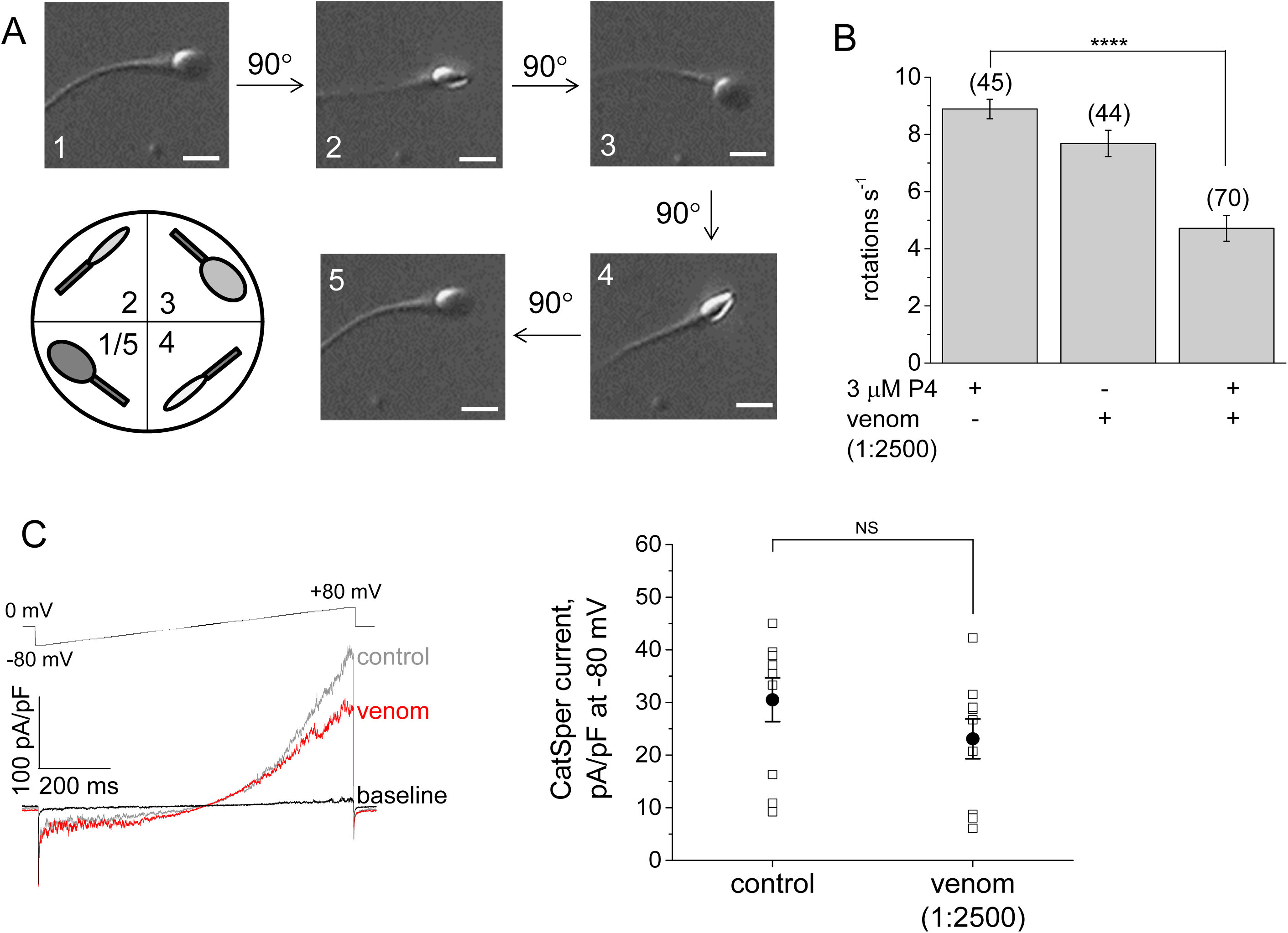
Sperm rotation significantly decreases when CatSper is activated but Hv1 is blocked. (A) Shown are representative snapshots of human capacitated spermatozoa performing rotational motion as they turn by 90 degrees within 15 ms timeframe; scale bar is 5 µm. The diagram below shows the position of the sperm head which is either parallel to the recording plane (1/5 and 3) or perpendicular (2, 4) to it. We have recorded sperm rotation along the long axis based on the changes in the reflected light intensity and changes in the head position as indicated in the diagram (see also Supplementary Movies 1-3). (B) The number of sperm rotations per second produced by human capacitated spermatozoa under control conditions and when subjected to the indicated treatment. ****P < 0.0001 *(calculated via t-test)*. The experiment was repeated three times with cells from two different human donors. (C) Venom does not affect human inward CatSper currents. Shown are representative CatSper currents in the absence (control, grey) and in the presence (venom, red) of the venom (same 1:2500 dilution). Right panel shows combined data recorded from 9 individual cells from three different human donors; NS-non-significant. Data are presented as the mean ± S.E.M. and (n) indicates number of individual cells analyzed.

### Hv1 is organized in asymmetrically positioned bilateral longitudinal lines

As Hv1 is an integral plasma membrane protein, the distance between the bilateral lines of ∼230 nm suggests that the Hv1 proteins are sequestered to one side of the flagellar midline. To examine this possibility, we performed 2-color 3D STORM imaging of both human Hv1 and the plasma membrane, using CM-DiI, a fixable variant of the lipophilic membrane dye DiI (DiIC_18_) (Shim et al., 2012) to label the latter (Fig. 2*B*). Although inhomogeneous, patchy staining was noted for the membrane stain, a phenomenon also observed in other cell types (Shim et al., 2012) and (Pan et al., 2018), the overall structure of the membrane is still well represented, as evidenced by the hollow cross-section. Remarkably, the bilateral lines of Hv1 were positioned asymmetrically to one side of the membrane, and this asymmetric distribution is maintained along the whole length of the sperm flagellum (Fig. S2 and S5A). Fig S2 demonstrates a situation where the sperm flagellum is fixed with varying degrees of rotation along its long axis. Such images provided a valuable observation that allows the tracing of bilateral lines along the whole length of the flagellum, and confirm that the two-line feature is well-maintained through the length of the principal piece (Fig. S2, and Fig. S5A) even as it rotates. Such asymmetric localization of Hv1 suggests a potentially important regulatory role. As previously mentioned, Hv1 activity can lead to flagellar alkalinization and thus create conditions for the activation of the pH-sensitive calcium channel CatSper. Calcium influx via CatSper triggers hyperactivation (Carlson et al., 2005) -an asymmetrical bending of the sperm flagellum (Fig. S6). In addition, CatSper activity is required for rheotaxis, which is a concerted flagellar motion of hyperactivated bends and rotation (Miki and Clapham, 2013). While CatSper is organized in a symmetric, quadrilateral arrangement along the sperm tail (Chung et al., 2017; Chung et al., 2014), Hv1, on the other hand, is positioned asymmetrically, thus suggesting a possible explanation for the specific tail motion.

### Hv1 inhibition decreases sperm rotation along the long axis

In order to reveal the physiological role Hv1 plays in sperm motility, the spermatozoa were capacitated in the presence of an Hv1 inhibitor-a hanatoxin containing venom from *Grammastola rosea*. As reported previously, such treatment diminishes Hv1 currents in human spermatozoa by 50% (Lishko et al., 2010) and Figure S8C. Hanatoxin is a known inhibitor of voltage-gated channels, and the only channel with strong voltage-gated characteristics functionally characterized in human sperm to date is Hv1. As expected, the treatment with venom did not alter inward currents via CatSper channel (Figure 3C), and no significant changes in sperm hyperactivation were observed. Next, sperm motility was recorded in the presence of progesterone and/or venom and compared with motility displayed by untreated spermatozoa. Full 360-degree rotation (Figure 3A-B) was significantly decreased when spermatozoa were treated with a combination of venom and progesterone (Figure 3A-B, Supplementary Movies 1-3). We were not able to achieve a complete inhibition of rotational movement, due to the fact the 1:2500 venom dilution only inhibits Hv1 by 50% (*24*). Interestingly, venom-treated sperm cells not only rotated less, but often displayed “partial” 180-degree rotation by flipping their heads from left to right (Supplementary Movie 3), similar to the motion produced by murine sperm (3, 26). It must be also noted that rodent spermatozoa lacked voltage-gated proton currents (Lishko et al., 2010) and Figure S8.

### The CatSper regulatory protein ABHD2 is organized in symmetrically positioned quadrilateral lines

Our next goal was to determine whether localization of the CatSper regulatory protein ABHD2 in human sperm resembles a CatSper or an Hv1-like distribution pattern. In human spermatozoa CatSper activation has two prerequisites: intraflagellar alkalinity and sperm exposure to extracellular progesterone. CatSper activation by progesterone occurs on a time scale of 100 ms (Lishko et al., 2011; Strünker et al., 2011), and is based on the elimination of the CatSper inhibitor 2-arachidonoylglycerol (2-AG) (Miller et al., 2016) by monoacylglycerol hydrolase ABHD2, which hydrolyses 2-AG in a progesterone-dependent manner. To enable the rapid progesterone-triggered activation of CatSper, it follows that the turnover of 2-AG should occur in close proximity to the channel itself. Therefore, we hypothesized that like CatSper in human and mouse sperm ((Chung et al., 2017; Chung et al., 2014) and (Fig. S3*D*)), its regulator ABHD2 follows a quadrilateral distribution and that the proton channel Hv1 is located in close proximity to the pair. With 3D STORM imaging, we assessed the localization of ABHD2 in the flagella of human sperm and found that this enzyme also follows a quadrilateral arrangement (Fig. 1 *C-D*). Superposition of single-color STORM images of ABHD2 (Figure 1*C-D*), Hv1 (Fig. 1*E-F*) and beta-tubulin (Fig. 1*G-H*) enabled a visual approximation of their relative distributions (Fig. 4*A-D*).

**Figure 4.**
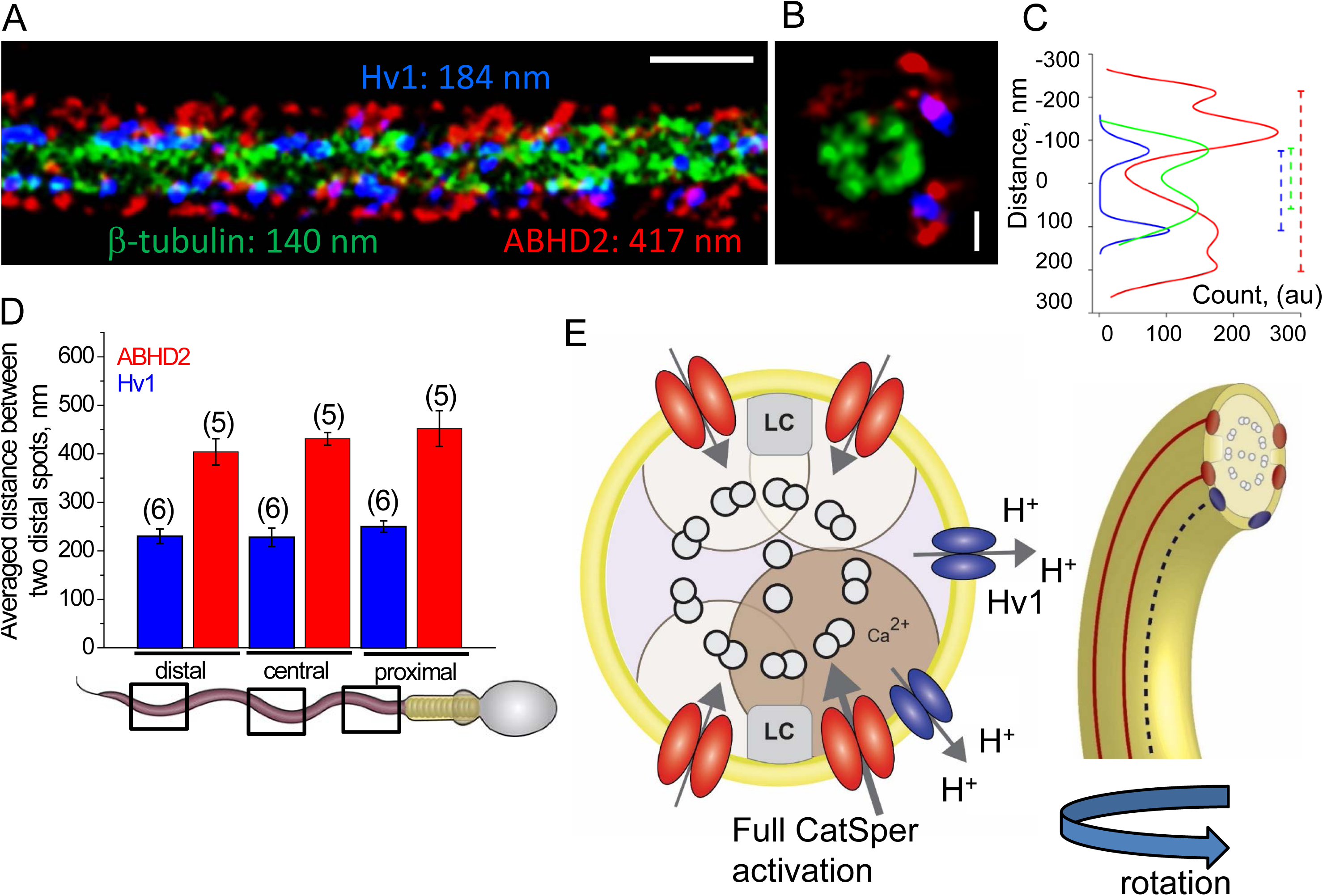
Proposed nanodomain structure of the flagellar controlled units in human sperm. (A) Superposition of panels (C), (E) and (G) from Figure 2 in x/y projections, as well as the relative widths of signals along the y-axis; scale bar is 500 nm. (B) Superposition of panels (D), (F) and (H) from Figure 2 in z/y projections. Scale bar is 100 nm. (C) Width distributions of targets as calculated from their localizations along the y-axis as shown in (A). (D) Averaged distance between Hv1 and ABHD2 spots at three regions along the human sperm flagellum as calculated from cross sections; (n) corresponds to the number of cross sections examined. Data are means +/-S.E.M. (E) The model suggests specific localization of CatSper/ABHD2 with respect to Hv1. The model is based on quadrilateral CatSper arrangement (red) in relation to asymmetrical Hv1 lines (blue). Proton extrusion resulted from Hv1 activity will lead to asymmetrical flagellar alkalinization and activation of only a subset of all flagellar CatSper channels, that are located in the close proximity to Hv1. This could lead to a flagellar rotation.

### Immunogold labeling detects Hv1 in the same compartment as shown by STORM imaging

To further elucidate the localization of Hv1 in the principal piece of human spermatozoa, immunogold labeling and transmission electron microscopy were performed (TEM; Fig. S5*B-E*). The majority of immunogold particles corresponding to Hv1 were found at the fibrous sheath in cross and longitudinal sections (FS), while significantly fewer particles were found at outer dense fibers (ODF) or microtubules (MT; Fig. S5*B-C*). The fibrous sheath is a unique cytoskeletal structure of the sperm principal piece that consists of two longitudinal columns (LCs; Fig. 1*B*) connected by semicircular ribs (Eddy et al., 2003; Fawcett, 1981). It surrounds the ODFs (Fig. 1*B*) and the sperm axoneme, a cytoskeletal structure comprised of microtubule doublets (Fig. 1*A-B*) arranged in a classical “9×2 + 2” pattern. ODFs are sperm tail-specific cytoskeletal structures, which support tail bending (Lindemann, 1996). In general, there are nine ODFs (Figure 1*B* and 4*D*) in the sperm tail. However, in the principal piece, two ODFs are replaced by inward projections of the LCs, which divides this part of the sperm tail into two planes – one with three ODFs (3 plane) and one with four ODFs (4 plane, Fig. 4*D*). The LC plane provides the flagellum with rigidity, so that the bending of the tail occurs perpendicular to this plane(Fawcett, 1981). Due to the thin (less than 70 nm thick) sections required for TEM and the sparse gold distribution along the sperm tail, we could not observe two immunogold dots in the same cross-section. Therefore, we analyzed immunogold distribution in the fibrous sheath (FS) of individual cross sections. Gold particles were predominantly found at the 3-plane of the principal piece, whereas a minor fraction accumulated at the 4-plane (Fig. S5D*-E*). Furthermore, gold particles were found preferentially at two distinct clusters at the 3 plane (position A and B, with 16 and 6 particles correspondingly; Fig. S5*D*). The distance between centers of the clusters “A” and “B” was 172 +/-5 nm (n= 24), which is close to the distance between Hv1 lines observed above with super resolution microscopy (Figs. 1*A*, 3*D* and Fig. S5*A*). It is quite possible that “AB” cluster could be positioned either on the left or the right side of the 3-plane, however further co-localization studies are needed to verify the exact localization of the cluster in relation to the chirality of the plane. These results confirm that Hv1 is distributed asymmetrically in the principal piece of human sperm, while CatSper channels cluster symmetrically on either side of the LC (Chung et al., 2014). Thus, it is therefore likely that proton efflux via Hv1 produces a unilateral alkalization of the flagellum and this results in the activation of only a subset of all CatSper channels, thus triggering specific flagellar motion.

### Simulation of proton efflux via Hv1 and its effect on the change of intraflagellar pH

Unilateral alkalization of the sperm intraflagellar environment has two prerequisites: 1) a significant local pH elevation within the Hv1 channel mouth, and 2) little or no impact on the distant CatSper channels. We tested the second requirement *in silico* by estimating the global pH change in a sperm tail from the action of all Hv1 channels operating simultaneously, both under the conditions of our experiments and under normal physiological conditions. The current corresponding to SI Appendix, Fig. S1*A* (right panel) in human sperm at +80 mV was fit to an exponential rising with a time constant of 0.51 s (Lishko et al., 2010) to a level of 28.2 pA lasting 3 s and assumed to flow uniformly across the surface of a cylinder, representing a human sperm tail at the principal piece region containing MES buffer. Fig. S7*A* shows that this would elevate submembrane pH substantially from 6.0 to 7.8, while average pH and pH at the axonemal core (core pH) would rise to 7.3. Under these conditions, the core and average pH rise sufficiently to significantly activate the rows of CatSper channels (Lishko et al., 2011) opposite and orthogonal to the row of Hv1 channels. However, such a stimulus is exceptional in both strength and duration, and does not reflect physiological conditions. Therefore, it was important to estimate what is likely to occur under physiological conditions between successive flagellar flicks, assuming a starting pH of 6.0 and a temperature of 37 °C. Fig. S7*B* shows pH now rising only to 6.006 in the time between flagellar flicks, clearly insufficient to influence distant CatSper channels. Note that the simulations overestimate pH rise by ignoring restorative processes such as hydrogen pumps, organelle actions, and metabolic processes.

Next, we estimated the steady-state level of pH that might be reached in the neighborhood of a single Hv1 dimer in the presence of natural buffers. Fig. S7*C* shows that hydrogen ions are depleted within 4 nm of the Hv1 channel “sink”, so that the pH rises to >7.4 within 4.5 nm from the “sink”. At 5 nm from “sink” the pH rises to 6.9, while only to 6.3 at 8-nm away. Under presumed natural conditions at 37 °C with more rapid hydronium diffusion and a native mobile buffer replacing MES, the local pH changes are somewhat less. However, local viscosity and tortuosity would raise the local pH even higher. These increases are extremely localized, and a CatSper target within about 3-5 nm of the “sink” would be strongly activated by an Hv1 dimer opening (Fig. S7*D*).

## Discussion

### Asymmetrical organization of the Hv1 channel is responsible for sperm rotation

On their route toward the egg, sperm cells use mostly symmetrical or snake-like tail motion. However, to overcome the high viscosity of the fallopian tubes or to detach from the tubal ciliated epithelium (Ho et al., 2009), spermatozoa must enhance their flagellar beating. This is achieved by hyperactivation – an asymmetrical flagellar bending (Fig. S6) that is triggered by calcium influx via the CatSper channel (Carlson et al., 2003; Ho et al., 2009) – and rotation (Ishijima et al., 1992; Kantsler et al., 2014; Miki and Clapham, 2013). Both motility types are required for rheotaxis, which allows sperm to overcome upstream fluid flow (Ishijima et al., 1992; Kantsler et al., 2014; Miki and Clapham, 2013). The asymmetrically positioned proton extrusion channels -Hv1 -could enhance rotation by producing local alkalinization in a close proximity to a subset of CatSper channels (Figure 4E). Hv1 therefore has the capacity to fully activate only a subset of CatSper channels, thus resulting in asymmetrical local calcium influx in the tail, and ultimately, creates an asymmetry in axonemal rigidity. Indeed, by using STORM imaging we found that Hv1 distribution in human flagellum follows this prediction (Figs. 2 and 4). While CatSper rows that are positioned opposite and orthogonal to the row of Hv1 channels still likely produce calcium influx, their net calcium influx should be smaller compared to that produced by CatSper in close proximity to Hv1 rows.

The human sperm flagellum has the largest Hv1 current density of all other cell types (Lishko et al., 2010). In fact, Hv1 current is the only proton current across the sperm plasma membrane that is detected by the patch-clamp technique. The concentration of a significant number of proton channels in such a small cellular domain as a flagellum in the close proximity to the pH-sensitive Ca^2+^ channel CatSper likely affects sperm physiology in a profound way. While the direct measurement of the intracellular pH changes due to sperm Hv1 activity has not been done yet, the fact that Hv1 has a powerful ability to deplete protons in its vicinity has been recently shown (De-la-Rosa et al., 2016). We have confirmed this finding by *in silico* estimation of the steady-state level of pH that might be reached in the neighborhood of a single Hv1 dimer. In fact, pH rises to 7.4 at 4.5 nm from the Hv1 “sink”, while only to 6.3 at 8-nm away. With these extremely localized pH increases, a CatSper channel complex or its putative pH-sensitive cytoplasmic region (Navarro et al., 2008) must be positioned within 5 nm of the Hv1 proton “sink” to be strongly activated by an Hv1. These results support our finding and explain why Hv1 needs to be positioned close to CatSper – to ensure local alkalinity, just enough to upregulate only a portion of CatSper channels. Given the fact that Hv1 can significantly change pH locally (De-la-Rosa et al., 2016), is expressed in human sperm at high density, and is physiologically active (Lishko et al., 2010), Hv1 can produce a significant local alkaline shift. However, such pH changes will occur in nanodomains that are localized only on one side of the flagellum and are likely active only while sperm cell moves and rotates. Since human sperm are quite fast movers (beat frequency of 24 Hz), it will require a powerful high-speed recording technique with sufficient sensitivity and spatial resolution to detect fluorescence changes in nanodomains. Such a limitation explains why Hv1-driven intracellular pH changes have not been yet detected in whole flagellar pH imaging.

Sperm Hv1 currents increase concurrently with sperm capacitation, as reported earlier (Lishko et al., 2010), further illustrating its relevance to the process of sperm activation in the female reproductive tract. Spermatozoa that swim in a linear trajectory usually produce smaller proton currents, while highly hyperactivated cells (based on their nonlinear swimming trajectory) produce upregulated Hv1 currents (Lishko et al., 2010). This indicates that nonlinear motility consistent with capacitation is a result of asymmetrical bending of the sperm flagellum and rotation, and is regulated by one or more factors which are asymmetrically localized.

Combining sperm electrophysiology and STORM imaging could also be useful for the approximate determination of single channel conductance, as current experimental limitations do not allow for the direct measurement of this ion channel characteristic. In the case of Hv1, we observed ∼ 7-8 Hv1 “clusters” per 1 µm of the flagellum in each longitudinal line (Fig. 3*B*). Since the length of the human principal piece is 40 µm, one can expect ∼320 “clusters” per line, which would result in 640 “clusters” per cell. According to the whole-cell recordings, Hv1 current density of human sperm is ∼ 25 pA/pF at +80 mV (Fig. S1*B*), which gives a permeability of 312.5 pS. Assuming that the open probability of Hv1 under such conditions is close to 100%, each cluster should have a permeability of ∼ 490 fS. Predicted single-channel permeability for Hv1 under these conditions is ∼240 fS (Cherny et al., 2003). Therefore, it is likely that each flagellar “cluster” is comprised of an Hv1 dimer (Tombola et al., 2008).

In conclusion, our results indicate that Hv1 localization in human sperm differs from that of ABHD2 and CatSper and is represented by off-centered bilateral longitudinal lines. This distribution may provide an explanation for sperm rotation along the long axis during sperm rheotaxis. Off-centered positioned Hv1 may selectively and unilaterally alkalinize only sub portion of the axoneme, which in turn activates a subset of CatSper channel clusters (Fig. 4*E*). Such asymmetric activation of sperm control units will result in a calcium increase in the portion of the flagellum, arrest dynein movement unilaterally and provide asymmetrical rigidity to the axoneme. This event can fuel a flagellar rotation as the means to relive the tension. Interestingly, sperm rotation is not dependent on the presence of the head, as observed by rotation displayed by decapitated human sperm flagellum (Supplementary Movie 4). Our results indicate that flagellar control units comprised of ABHD2, CatSper, and Hv1 are specifically positioned in order to ensure fast signal transduction.

## Materials and Methods are provided in the extended online materials

## Acknowledgements

This work was supported by NIH Grants R01GM111802 and R21HD081403, Pew Biomedical Scholars Award 00028642, Alfred P. Sloan Award FR-2015-65398, and Packer Wentz Endowment Will (to P.V.L.). K.X. is a Chan Zuckerberg Biohub investigator and acknowledges support from the Pew Biomedical Scholars Award and NSF under CHE-1554717. We thank Jean-Ju Chung for sharing anti-CatSper delta antibody and Reena Zalpuri and Kent McDonald from UC Berkeley EM facility for help with TEM images acquisition. We also thank James F Smith (UCSF, Urology Department) for sharing samples of the human epididymal fluids, obtained with patient consent.

## Author Contributions

M.R.M. and P.V.L. conceived the project, designed the experiments. M.R.M., S.J.K., K.X. and P.V.L. wrote the manuscript. M.R.M. performed immunostaining and electrophysiology. P.V.L. performed pilot electrophysiology experiments. S.J.K., M.W., and K.X. performed STORM and SEM sample preparation, image acquisition, and data analyses. S.A.M. helped with pilot electrophysiology experiments from animal sperm cells. N.M. performed immunoelectron staining and sperm motility analysis. S.M. helped with sperm motility acquisition and analysis. R.S.Z. performed in silico measurements, analyses and commented on the manuscript. P.V.L. and K. X. supervised and led the project. All authors discussed the results and commented on the manuscript.

---

**Figure S1.**
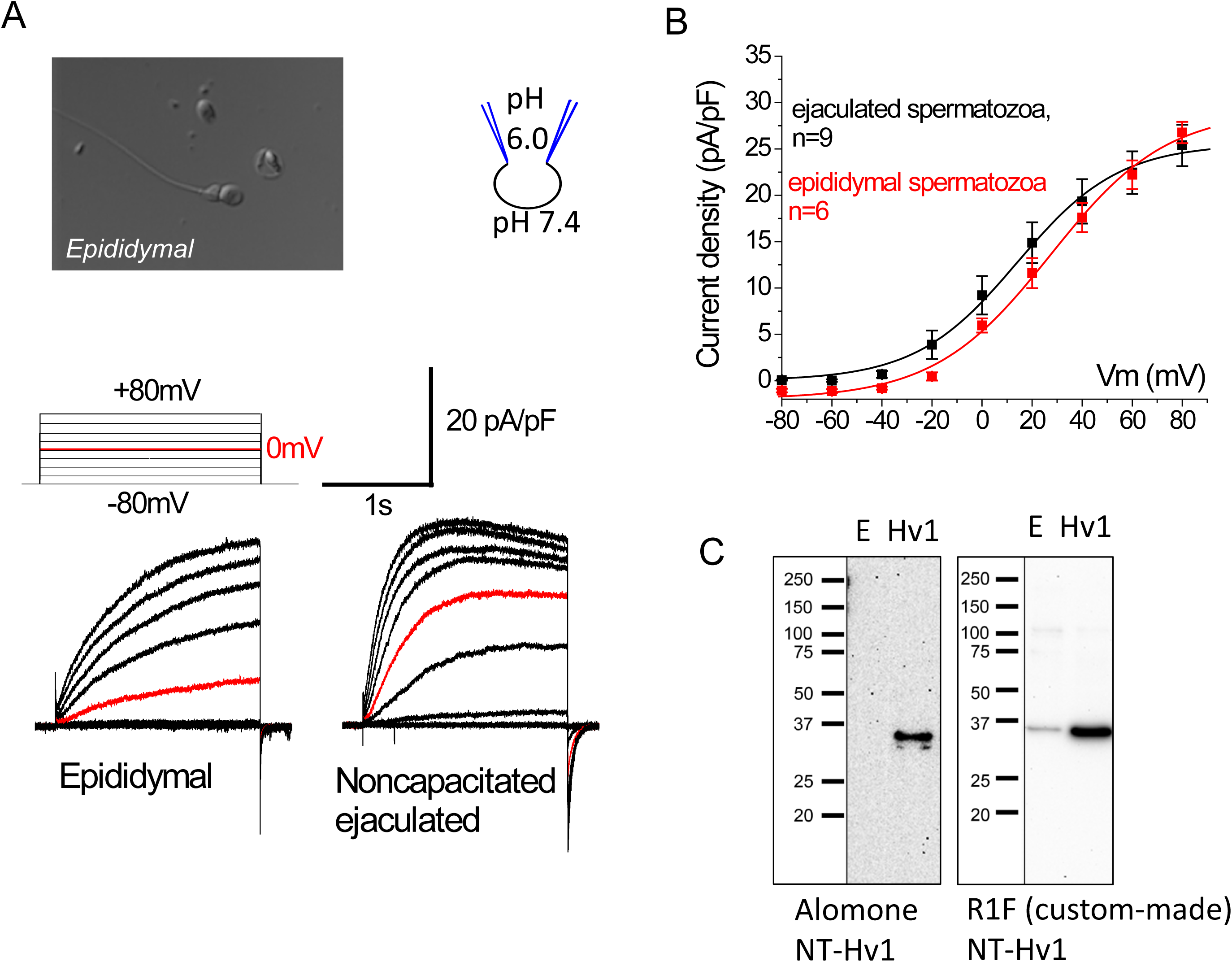

**Figure S2.**
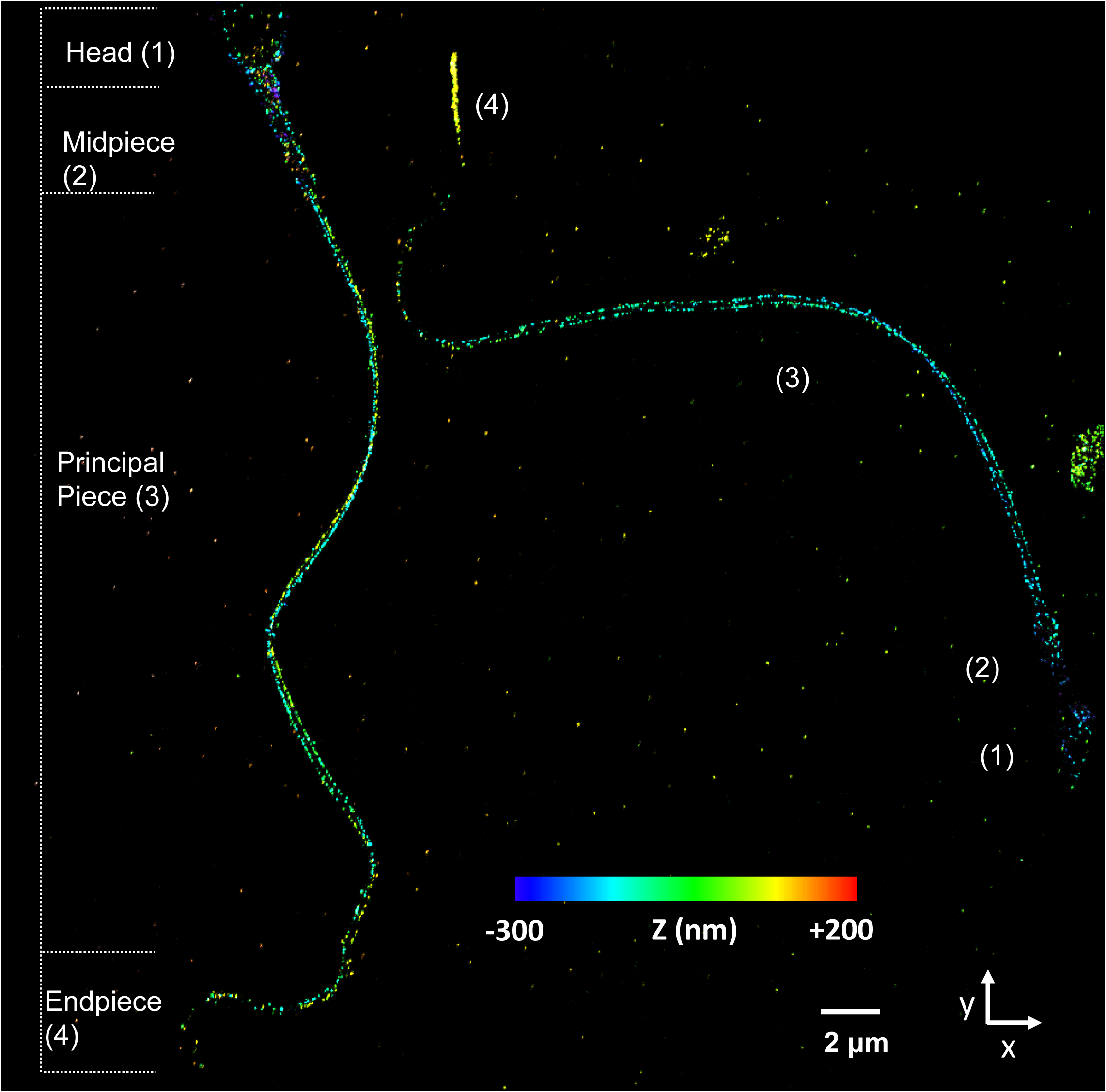

**Figure S3.**
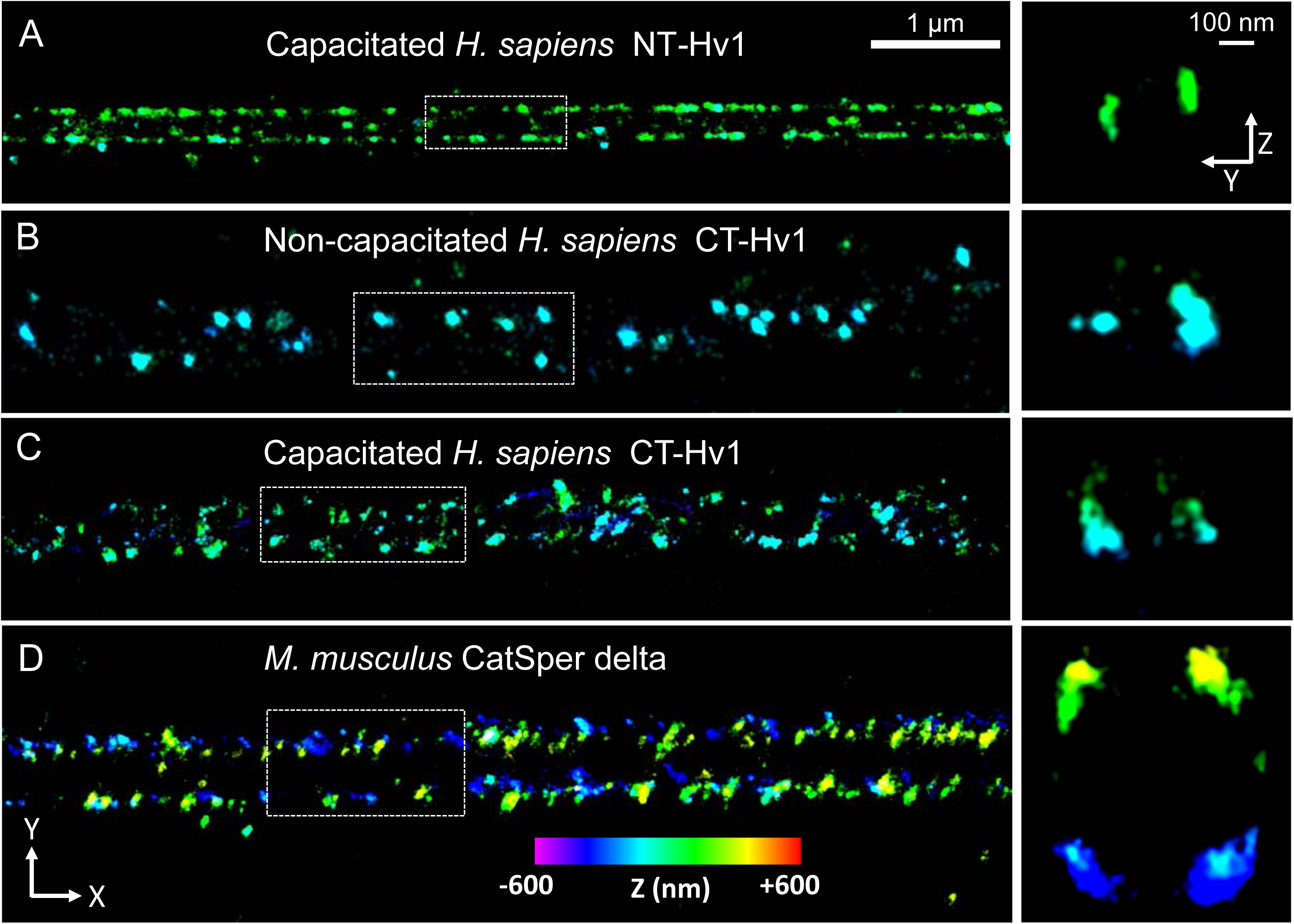

**Figure S4.**
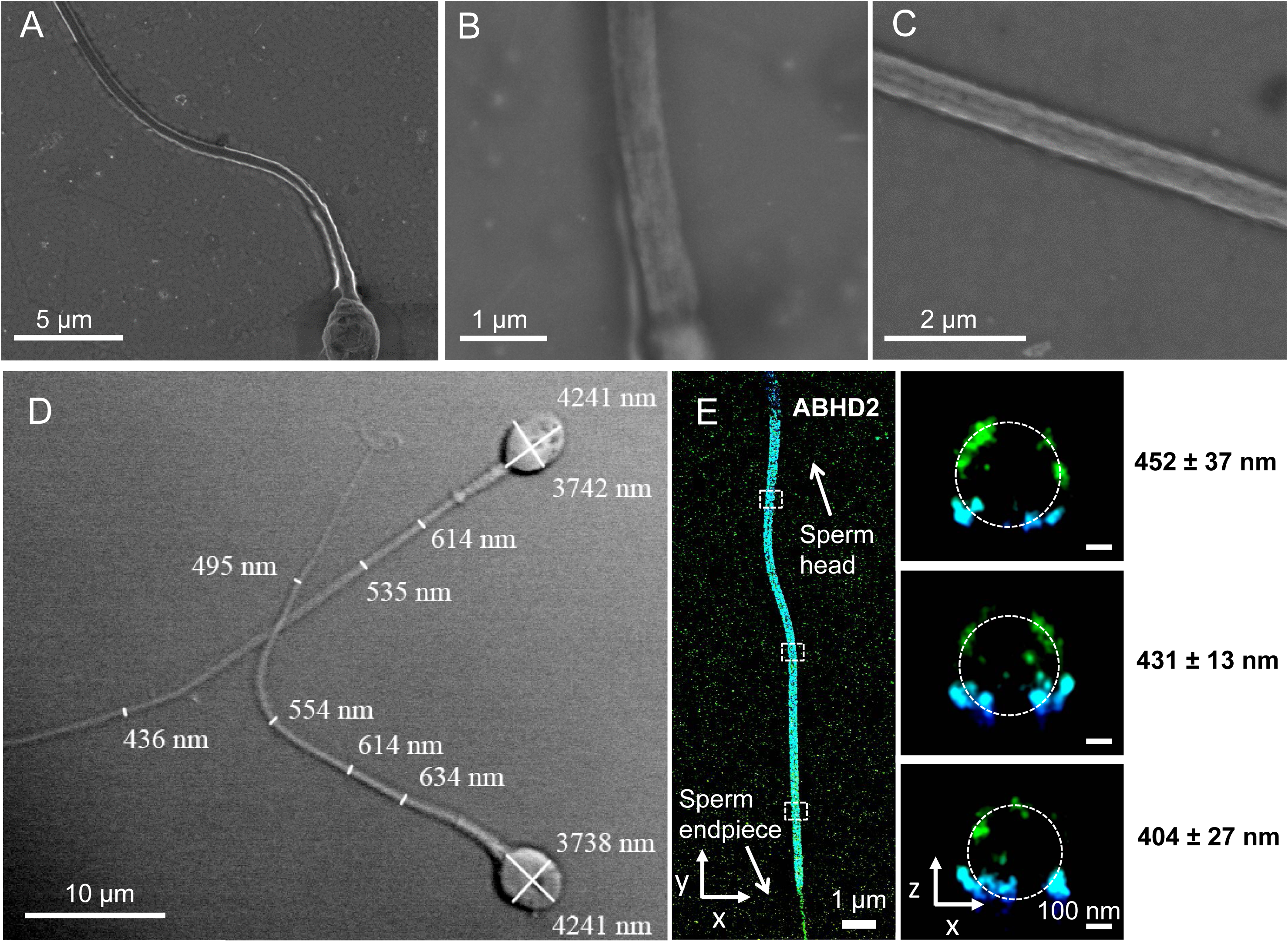

**Figure S5.**
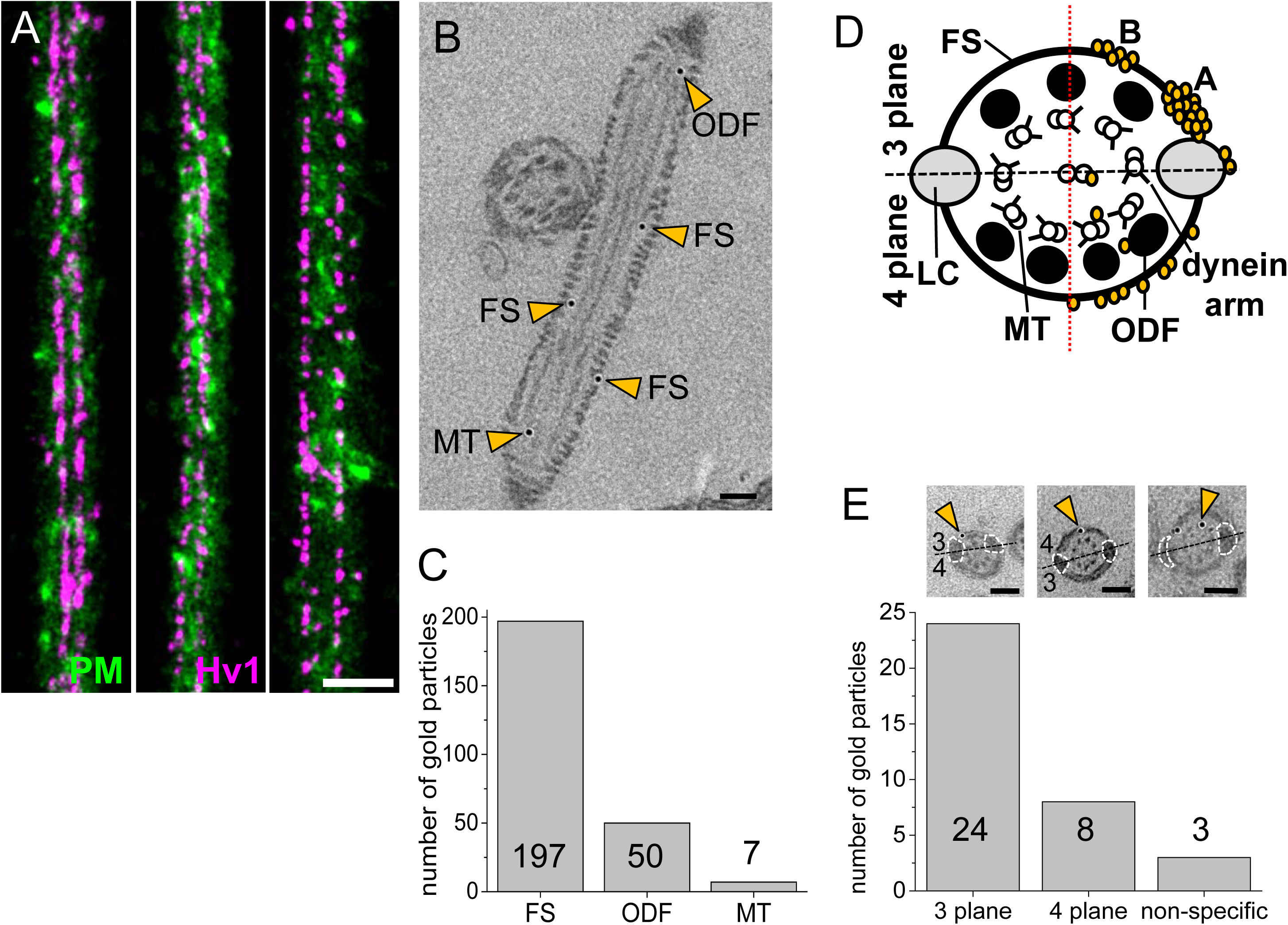

**Figure S6.**
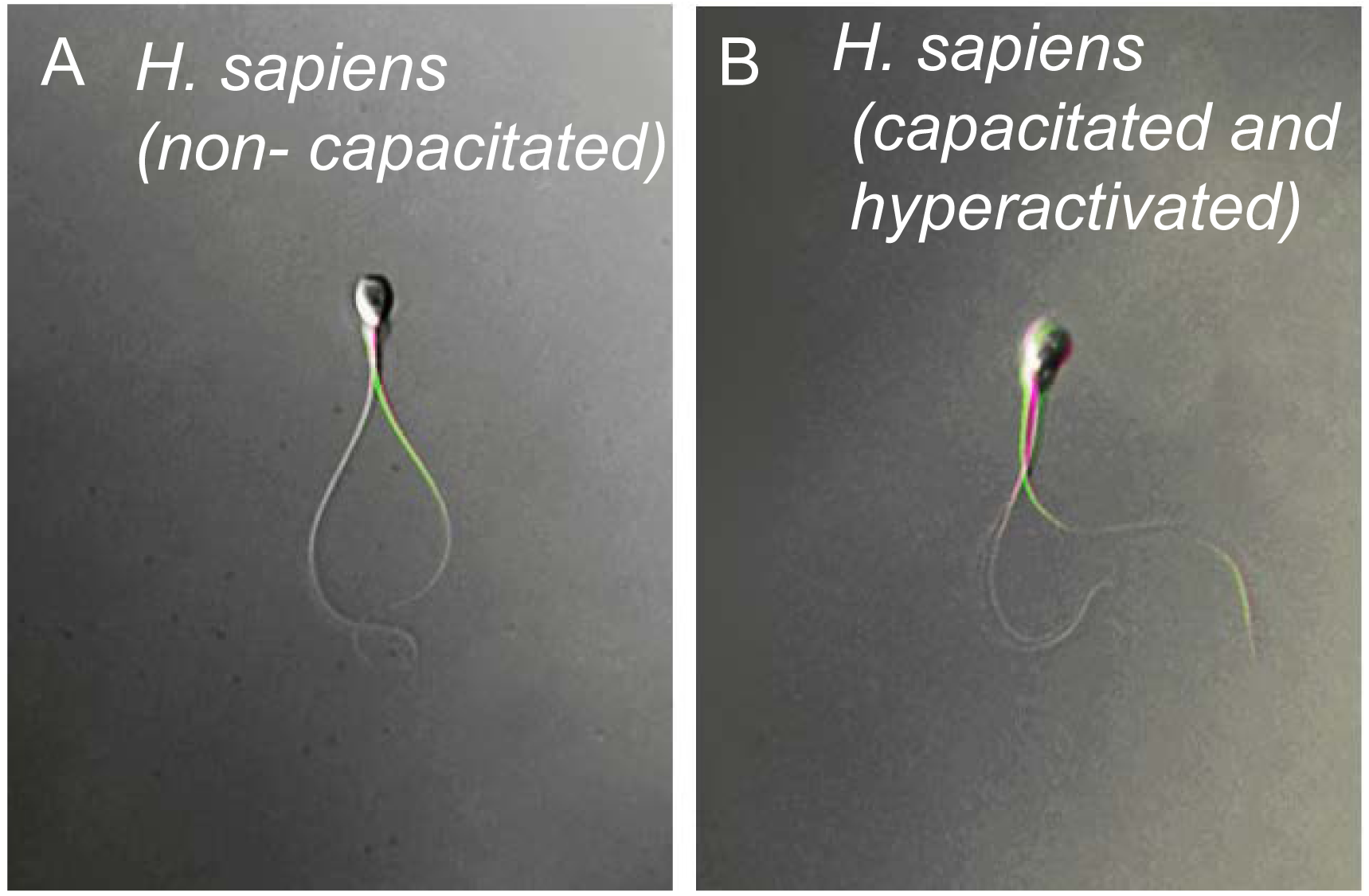

**Figure S7.**
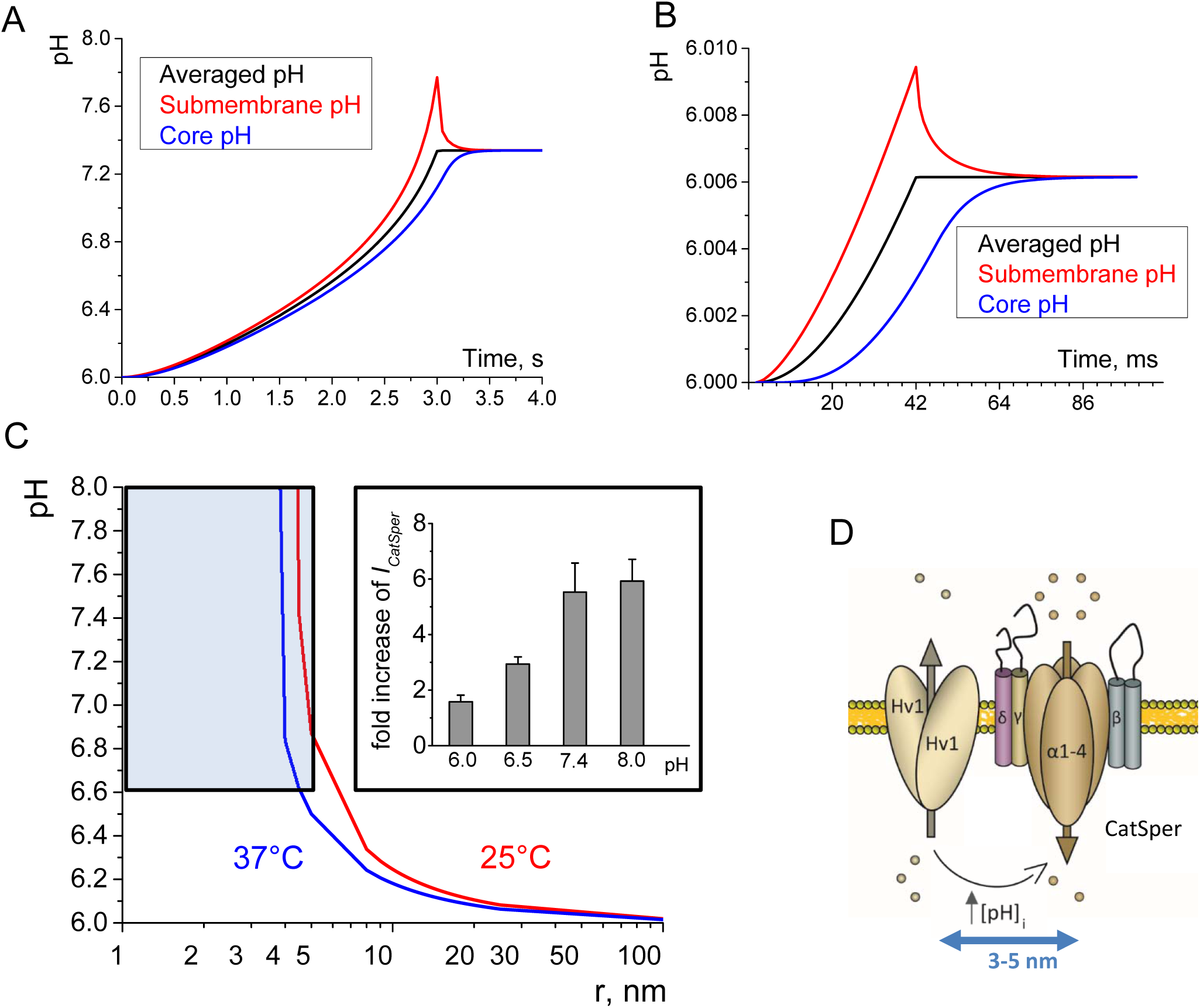

**Figure S8.**
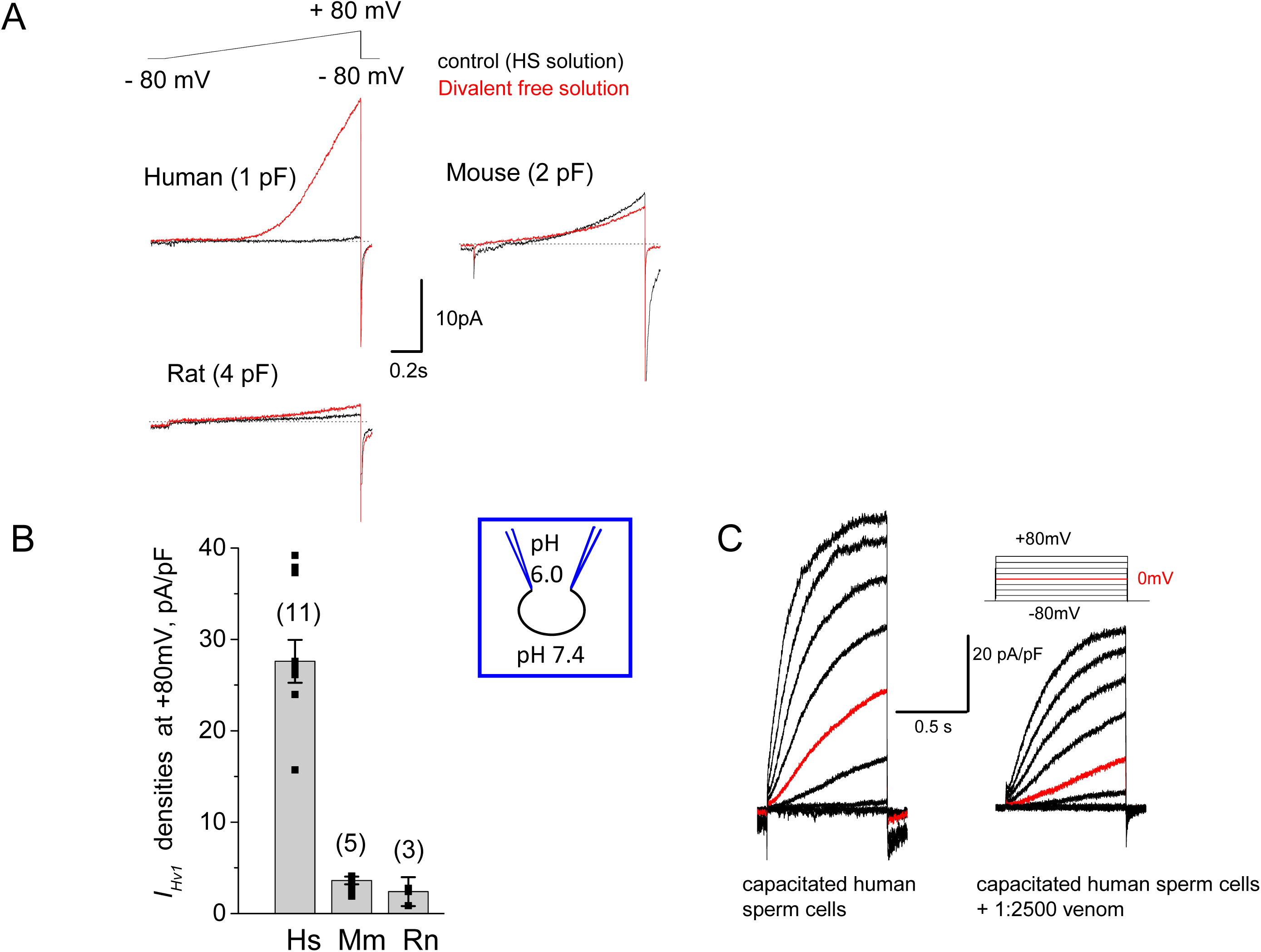

