## Supplementary Materials for "Asymmetrically Positioned Flagellar Control Units Regulate Human Sperm Rotation"

**SUPPORTING INFORMATION**
**SI Appendix Figure legends:**

**Fig. S1. Hv1 is present in human epidydimal and ejaculated spermatozoa.** (A) Representative recordings from a human noncapacitated ejaculated spermatozoon and a sperm cell isolated from human *cauda epididymis*. Traces were recorded in response to voltage steps from the holding potential of -80mV with increasing voltage steps (20 mV increment) up to +80 mV. The traces recorded at 0 mV are shown in red. The insert shows a picture of human epididymal sperm cells. The diagram depicts a recording pipette with the cell attached and indicates intracellular pH and pH in the bath. (B) Averaged current density shows human sperm Hv1 current in both sperm preparations. Data are means +/- S.E.M, and n indicates number of sperm cells tested. (C) Representative western blots showing CHO cell lysates from cells transiently transfected with empty vector (E), or expression vector bearing human Hv1 (Hv1). Both primary antibodies were used in 1:10,000 dilution. The blots were probed with a commercially available antibody from Alomone Labs (Alomone NT-Hv1) and an in-house custom-generated anti-Hv1 antibody (R1F NT-Hv1, reported in (Lishko et al., 2010))

**Figure S2. STORM imaging of Hv1 in human sperm cells.** Shown are two human sperm cells immunostained for Hv1. The color reflects the relative distance from the focal plane along the z axis as shown with the color scale bar. Sperm regions (the head, midpiece, principal piece and the endpiece are indicated).

**Figure S3. STORM imaging of the human and mouse sperm flagellum.** (A) Left panel: immunostaining of a human capacitated spermatozoon probed with anti-Hv1 antibody (Alomone, N-terminal antigen). Right panel: the cross section from the same flagellum (from the boxed region) shows double-line Hv1 staining. (B) Left panel: immunostaining of a human noncapacitated spermatozoon probed with anti-Hv1 antibody (Sigma, C-terminal antigen). Right panel: the cross section from the same flagellum (from the boxed region) shows double-line Hv1 staining. (C) Left panel: immunostaining of a human capacitated spermatozoon probed with anti-Hv1 antibody (Sigma, C-terminal antigen). Right panel: the cross section from the same flagellum (from the boxed region) shows unchanged double-line Hv1 staining. (D) Left panel: immunostaining of a mouse caudal spermatozoon probed with anti-CatSper antibody. Right panel: the cross section from the same flagellum (from the boxed region) shows quadrilateral CatSper arrangement. Left panels in (A-D) show conventional x/y projections of the sperm flagellum, while right panels show cross-sections in y/z projections from the corresponding cells. Color reflects the relative distance from the focal plane along the z axis as shown on the color scale bar in (D). Scale bars on x/y projections are 1 μm, while scale bars on y/z projections are 100 nm.

**Figure S4. Scanning electron microscopy (SEM), differential interference contrast (DIC) and STORM imaging of ABHD2 to estimate the average flagellar diameter of the human sperm tail.** (A-C) Representative images of several human sperm revealed by SEM of graphene-protected, non-dehydrated samples. (D) Representative DIC image of human spermatozoa. (E) STORM imaging of the human sperm flagellum probed with anti-ABHD2 antibody as shown in Figure 2c. Cross sections (right panels) were taken from corresponding regions of the x/y projection (left panel) as indicated by dotted boxes; from top to bottom, ROIs were selected at distances of 5, 14, and 20 μm from the beginning of the principal piece (PP). Right panels show averaged distances between two distal ABHD2 spots acquired at the corresponding tail segments; data are means +/- S.E.M. with n=5. Scale bars are 1 μm (left); 100 nm (right). 3D color scale is as shown in Fig. 2.

**Figure S5. Immunoelectron microscopy of Hv1 localization is consistent with bilateral arrangement revealed by STORM.** (A) Two-color STORM images of Hv1 (magenta) and the plasma membrane (Dil, green) taken at three different parts of PP (from left to right: the distal, middle, and proximal PP parts). Note also the different apparent distances between the two Hv1 line patterns in this 2D projection image due to different local rotational orientations of the PP. Scale bar is 500 nm. (B) Gold particles are localized mainly to the fibrous sheath (FS), and to a lesser extend to outer dense fibers (ODFs) and to microtubules (MT) as shown in a representative longitudinal section through the principal piece of a human sperm cell; scale bar is 200 nm. (C) Gold particle quantification along longitudinal and cross sections shows that the majority of gold particles are detectable at the FS (197 particles), while fewer particles are found at the ODFs (50 particles) and MT (7 particles). (D) Cartoon illustrating gold particle distribution at individual positions combined from 35 individual cross sections. Since it is not possible to determine if gold particles were located on the left or on the right side of the sperm tail, all gold particles were plotted onto the right side. Gold particles are found predominantly on the "3 plane" forming two distinct clusters: cluster "A" (6 particles) and cluster "B" (16 particles). (E) Gold particle quantification of individual cross sections at the FS of the 3 plane (24 particles) and 4 plane (8 particles) as well as at unspecific regions (ODFs or MT; 3 particles); scale bars are 200 nm. For this analysis only PP cross sections with clearly identifiable LCs and ODFs were included.

**Figure S6. The maximal tail amplitudes in either direction of non-capacitated and capacitated human spermatozoa were superimposed to show different motility patterns.** (A) The two images of a human non-capacitated spermatozoon show symmetrical flagellar bend. (B) After capacitation, the flagellum bends asymmetrically resulting in a hyperactivated motility pattern. Each flagellum is shown in two different positions ~ 42 ms apart: one is colored and other is not.

**Figure S7. Simulated sperm intraflagellar pH changes caused by Hv1 activity.** (A) The total pH change in a whole sperm tail principal piece from the action of all Hv1 channels operating simultaneously, modeled by $H^+$ current passing uniformly across the surface of a cylinder representing a human sperm tail at the principal piece region: 0.5-μm-diameter and 40-μm-long. Sperm tail pH was calculated for three positions: submembrane (red; estimated as ~ 5 nm shell below the plasma membrane), core (at the center of a sperm axoneme, blue), and total volume-averaged (black). Parameters were chosen based on those in Fig. S1A (right panel), with a voltage clamp of +80 mV, while the channel stayed open for 3 s. (B) The effects of $H^+$ efflux under physiological conditions, with a 0 mV membrane potential and activation time of 40 ms, corresponding to the duration between flagellar flicks. Note that while proton efflux was simulated uniformly for simplicity, efflux along asymmetrical lines of Hv1 channels would probably cause alkalinization and affect distant CatSper channels in a way similar to the "core pH" (blue line). No pH restorative processes are included. (C) $H^+$ efflux from an Hv1 dimer is represented as a point sink from a semi-infinite space opening at time zero. The local steady-state pH changes near the dimer are plotted for conditions corresponding to the experiments at 25°C (red) and physiological conditions at 37°C (blue). The inset (extracted from data shown in Figure 2d from (Lishko et al., 2011)) plots the fold increase in CatSper inward current activated by 500 nM progesterone in response to CatSper activity in the absence of the progesterone at different intraflagellar pHs. A significant effect begins above pH 6.5, and this region is indicated by a blue square. (D) A separation of a few nm (up to 5 nm) between Hv1 and CatSper channels in overlapping rows would fall within this region and allow CatSper to be directly regulated by Hv1. All simulations include a native fixed buffer, plus a native mobile buffer in (B) and (C, blue), which were replaced by 100mM MES in (A) and (C, red).

**Figure S7. The presence of the functional proton channel Hv1 in mammalian sperm.** (A) Representative Hv1 current recording from sperm cells of different species as indicated. (B) Averaged currents ($I_{Hv1}$) density recordings from sperm of 3 different species; (n)- indicates the number of experiments. Data are means +/- S.E.M. Abbreviations are: noncapacitated human (Hs; *Homo sapiens*), mouse (Mm; *Mus musculus*), rat (Rn; *Rattus norvegicus*). (C) Representative Hv1 current recordings from a human capacitated ejaculated spermatozoon and sperm cell capacitated in the presence of tarantula venom. Traces were recorded in response to voltage steps from the holding potential of -80mV with increasing voltage steps (20 mV increment) up to +80 mV. The traces recorded at 0 mV are shown in red. The diagram depicts a recording pipette with the cell attached and indicates intracellular pH and pH of the bath solution.

**Supplementary Movie 1. Capacitated human spermatozoa in the presence of progesterone.** Representative cells are shown. Cells were capacitated for 4.5 hours and treated with progesterone immediately before recording. Motility was recorded using 1000 fps protocol and slowed down for playback with 200 fps.

**Supplementary Movie 2. Capacitated human spermatozoa in the presence of venom diluted 1:2500.** Representative cells are shown. Cells were capacitated for 4.5 hours with venom. Motility was recorded using 1000 fps protocol and slowed down for playback with 200 fps.

**Supplementary Movie 3. Capacitated human spermatozoa in the presence of progesterone and venom (1:2500).** Representative cells are shown. Cells were capacitated for 4.5 hours with venom and treated with progesterone immediately before recording. Motility was recorded using 1000 fps protocol and slowed down for playback with 200 fps.

**Supplementary Movie 4. Capacitated human sperm flagellum in the presence of progesterone.** Representative flagellum of the decapitated sperm is shown. Cells were capacitated for 4.5 hours and treated with progesterone immediately before recording. Motility was recorded using 1000 fps protocol and slowed down for playback with 200 fps.

### SI Materials and Methods

**Animals.** Male C57BL/6 mice were purchased from Harlan Laboratories (Livermore, CA) and were kept in the Animal Facility of the University of California, Berkeley. All experiments were performed in accordance with NIH Guidelines for Animal Research and approved by UC Berkeley Animal Care and Use Committee under the approved protocol MAUP #R352-012. Animals were humanely euthanized according to ACUC guidelines, and sperm were collected as described previously(Wennemuth et al., 2003).

**Healthy donors and isolation of human ejaculated spermatozoa.** A total of 18 healthy volunteers aged 21-38 were recruited to this study and all experimental procedures utilizing human derived samples were approved by the Committee on Human Research at the University of California, Berkeley, protocol number 2013-06-5395. Freshly ejaculated semen samples were obtained by masturbation and spermatozoa purified by the room temperature swim-up technique as described (Lishko et al., 2011) using artificial human tubal fluid solution (HTF, in mM): 98 NaCl, 4.7 KCl, 0.3 $KH_2PO_4$, 2 $CaCl_2$, 0.2 $MgSO_4$, 21 HEPES, 3 glucose, 21 lactic acid, 0.3 sodium pyruvate, pH 7.4 (adjusted with NaOH).

**Electrophysiology.** All recordings were performed as described in (Lishko et al., 2013; Lishko et al., 2011). Briefly, gigaohm seals between patch pipette and human spermatozoa was formed at the cytoplasmic droplet. If the cytoplasmic droplet was inconspicuous, spermatozoa were patched at the neck region. Seals were formed in high saline (HS) solution containing: 130 mM NaCl, 5 mM KCl, 1 mM $MgSO_4$, 2 mM $CaCl_2$, 5 mM glucose, 1 mM sodium pyruvate, 10 mM lactic acid, 20 mM HEPES, pH 7.4 adjusted with NaOH, 320mOsm/L. Transition into the whole-cell mode was performed by applying light suction in combination with short voltage pulses. Access resistance was 25-40 MΩ. Cells were stimulated every 5 s. Data were sampled at 2-5 kHz and filtered at 1 kHz. For proton currents: pipettes for whole-cell patch-clamp recordings (20–30 MΩ) were filled with 135 mM N-methyl-D-glucamine (NMDG), 5 mM ethylene glycol tetraacetic acid (EGTA), and 100 mM MES and pH adjusted to 6.0 with methanesulfonic acid. Proton currents were recorded in divalent-free (DVF) bath solution comprising 130 mM NMDG, 100 mM HEPES, and 1 mM EDTA and pH adjusted to 7.4 with methanesulfonic acid. For monovalent CatSper recordings (Figure 3C) pipettes (11-17 MΩ) were filled with (in mM): 130 Cs-methanesulfonate, 70 HEPES, 3 EGTA, 2 EDTA, 0.5 TrisHCl, pH 7.4 adjusted with CsOH. Bath divalent-free (DVF) solution for recording of monovalent CatSper currents contained (in mM): 140 Cs-methanesulfonate, 40 HEPES, 1 EDTA, pH 7.4 adjusted with CsOH. All electrophysiology experiments were performed at ambient temperature and currents elicited by voltage ramps or step protocols as indicated for each individual experiment Data were analyzed with OriginPro 9.0 and Clampfit 9.2. Statistical data were calculated as the mean ± S.E.M., and (n) indicates number of experiments.

**STORM imaging.** Dye-labeled cell samples were mounted on glass coverslips with a standard STORM imaging buffer consisting of 5% (w/v) glucose, 100 mM cysteamine, 0.8 mg/mL glucose oxidase, and 40 μg/mL catalase in 1M Tris-HCl (pH 7.5)(Huang et al., 2008; Rust et al., 2006). Coverslips were sealed using Cytoseal 60. STORM imaging was performed on a homebuilt setup based on a modified Nikon Eclipse Ti-E inverted fluorescence microscope using a Nikon CFI Plan Apo λ 100x oil immersion objective (NA 1.45). Dye molecules were photoswitched to the dark state and imaged using either 647- or 560-nm lasers (MPB Communications); these lasers were passed through an acousto-optic tunable filter and introduced through an optical fiber into the back focal plane of the microscope and onto the sample at intensities of ~2 kW $cm^{-2}$. A translation stage was used to shift the laser beams towards the edge of the objective so that light reached the sample at incident angles slightly smaller than the critical angle of the glass-water interface. A 405-nm laser was used concurrently with either the 647- or 560-nm lasers to reactivate fluorophores into the emitting state. The power of the 405-nm laser (typical range 0-1 W cm-2) was adjusted during image acquisition so that at any given instant, only a small, optically resolvable fraction of the fluorophores in the sample were in the emitting state. Emission was recorded with an Andor iXon Ultra 897 EM-CCD camera at a framerate of 110 Hz, for a total of ~80,000 frames per image. For 3D STORM imaging, a cylindrical lens of focal length 1 m was inserted into the imaging path so that images of single molecules were elongated in opposite directions for molecules on the proximal and distal sides of the focal plane. Two-color imaging was performed via sequential imaging of targets labeled by Alexa Fluor 647 and CM-DiI. Nanoscale color channel alignment was achieved by adding 0.002% (w/v) yellow-green fluorescent beads into the STORM imaging buffer (FluoSpheres; Thermo F8811). Adherent beads near the cell to be imaged were activated and imaged in each color channel for the entire imaging duration. The resulting localizations were used to align the color channels in 3 dimensions.

**Reagents.** Progesterone was purchased from CalBiochem (EMD Millipore, Darmstadt, Germany). All other compounds were from Sigma-Aldrich (St. Louis, MO) unless otherwise specified. Primary antibodies: rabbit anti-ABHD2 IgG (C14214) was obtained from Assay Biotech (San Francisco, CA). Monoclonal mouse anti-beta-tubulin IgG1 (T5201) were purchased from Sigma-Aldrich. The anti-actin antibody (ab3280) was from Abcam (Cambridge, UK). Rabbit anti-Hv1 IgG (AHC-001; ALA, against N-terminus) was from Alomone Labs (Jerusalem, Israel), rabbit anti-Hv1 IgG (HPA039329; against C-terminus) were from Sigma, and affinity purified rabbit-anti-Hv1 (R1F, against N-terminus) IgG was custom-made purified (Lishko et al., 2010). The CatSper-delta antibody was received from Jean-Ju Chung (Chung et al., 2014). Anti-ABHD2 (C14214) were from One World Lab. Secondary antibodies: anti-rabbit AF647 or AF488; anti-mouse AF647; anti-mouse AF647 were from Invitrogen. Anti-rabbit CF680 and anti-rat CF680 were made by conjugating CF680 NHS-ester (Biotium) to unlabelled corresponding secondary antibodies (Jackson Immunoresearch). Human (NLH-06) and mouse (NLM-06) testis lysates were purchased from G-Biosciences (Geno Technology). *Grammostola rosea* venom was purchased from Spider Pharm (http://spiderpharm.com/).

**Isolation of human epididymal spermatozoa.** Men with proven fertility who were undergoing sperm retrieval procedures in the UCSF Center for Reproductive Health agreed to donate unused portions of surgical specimens. As part of the ongoing IRB-approved UCSF LIFE (Lifestyle, Fertility, and Evaluation) study, men enrolled in the study had a documented history of prior paternity and had undergone a vasectomy in the past. As part of routine clinical care, these men elected to undergo a sperm retrieval procedure (microscopic epididymal sperm aspiration) combined with in vitro fertilization (IVF) or a vasectomy reversal. Aliquots of epididymal fluids were used for the present study with patients' consents. Epididymal spermatozoa were isolated from the samples as described[29].

**In vitro capacitation.** The capacitation medium (HSB/BSA) consisted of HS solution supplemented with 15 mM $NaHCO_3$ and 5% BSA. Human sperm cells were capacitated for 4-5 h at 37°C and 5% $CO_2$ in HSB/BSA alone or in HSB/BSA spiked with either 3 μM P4, G. rosea crude venom (1:2500) or a combination of venom (1:2500) plus P4 (3 μM).

**Assessment of sperm rotation.** 100 μl of the sperm suspension were transferred to the recording chamber. Free-swimming sperm were examined on an inverted microscope (Olympus IX-71) equipped with a 60 ×/1.20 W objective. 500 ms-long movies were taken with a digital high-speed Memrecam GX-1 camera (NAC Image Technology, Simi Valley, CA) and collected with the Memrecam GXLink software, version 3.20 (NAC Image Technology) at 1,000 frames per second from a 640 × 480 pixel region of the camera chip and stored in AVI format. Movies were replayed with ImageJ version 1.44o (http://rsb.info.nih.gov/ij/) to assess the number of rotations of individual cells. Since each movie was only 500 ms long, values were multiplied by 2 to obtain rotations per second. Data were evaluated with the OriginPro 9.0 software (Origin Lab, Northampton, MA) and expressed as means ± SEM. Statistical significance (t-test) was indicated by: ****, $p < 0.0001$, ***, $p < 0.001$, **, $p < 0.005$ and *, $p < 0.05$. No variation between human donors were noticed, and control and venom- treated samples were donor matched for the analysis. It is important to mention that while control untreated sperm cells can engage in full 360-degree rotation (or rolling) movement, the venom treated cells only rotate half way: producing 180-degree rotation. We have analyzed each frame by looking at the "blinking" of the head as the indication of the rotatory movement, as well as by determining the entire head position to discriminate between a complete 360-degree rotation and a partial, 180-degree flipping. We have also determined rotation frequency of the full 360-degree rotating sperm only and ignored non-rotating sperm cells. In addition, we have asked three independent observers to analyze the movies in the unbiased way and all three respondents interpreted the movies correctly, pointing to the venom/P4 -treated cell as those that show decreased rotation.

**Immunocytochemistry.** Purified spermatozoa (concentration ~$10^7$ cells/ml) were plated onto coverslips in HS and allowed to attach for 10 min. The cells were fixed with 4% paraformaldehyde (PFA) in 1x PBS (Phosphate Buffered Saline) for 10 min and washed twice with PBS. Additional fixation was performed for samples not subsequently labeled with CM-DiI using 100% ice-cold methanol for 1 min with two washes in 1x PBS. Cells were permeabilized for 5 min with 0.25% Triton or 0.1% saponin in PBS followed by two PBS washes and blocked for one hour in PBS supplemented with 3% gamma-globulin free BSA (bovine serum albumin, fraction V). Immunostaining was performed in blocking solution. Detergent-treated cells were incubated with primary antibodies overnight at 4°C. After extensive washing in PBS, secondary antibodies were added for 60 min. The dilutions of primary antibodies were 1:100 and secondaries were 1:200. For membrane visualization, cells were then incubated with the lipophilic membrane dye CM-DiI (3H-Indolium, 5-[[[4-(chloromethyl)benzoyl]amino]methyl]-2-[3-(1,3-dihydro-3,3-dimethyl-1-octadecyl-2H-indol-2-ylidene)-1-propenyl]-3,3-dimethyl-1-octadecyl-, chloride, Invitrogen) at 5 μM in PBS for 20 min at room temperature. After 3 washes with PBS, cells meant for confocal imaging were mounted with ProLong Gold antifade with DAPI reagent (Life Technologies, Carlsbad, CA) or if cells were to be used for super-resolution imaging, coverslips were post-fixed for 10 min in 3.7% PFA and 0.1% glutaraldehyde, washed twice with PBS and stored in PBS until imaging.

**Image processing.** The raw STORM data were analyzed according to previously described methods (Huang et al., 2008; Rust et al., 2006). Briefly, optical astigmatism elongated images of single-molecules along either the vertical or horizontal axis for molecules above or below the focal plane, respectively. Intensity-based thresholding was used to detect single molecule blinking events, which were subsequently fit to elliptical 2D Gaussians. The resulting centroid positions were used to determine the superresolved lateral positions of each single molecule. Axial positions were determined by a calibration curve which mapped the ellipticity of each fitted Gaussian to its position above or below the focal plane. The calibration curve was generated by imaging and measuring the ellipticities of fluorescent beads adhered to a coverglass, while moving the sample smoothly through the focal plane (39). Final superresolution images used in figures were generated by projecting all localizations in the specified plane and blurring each point to a 2D Gaussian profile. The width of the Gaussian was chosen to match our resolution, which was experimentally determined by measuring the full width at half maximum in each dimension of a cluster of localizations from a single isolated fluorophore (38,39).

**Geometric characterization of STORM cross sections.** Distance measurements were obtained from STORM data using custom MATLAB routines. For a direct width comparison of different targets, 1D histograms were generated on straight, cropped sections of sperm tails for the coordinate perpendicular to the tail axis. Histogram bin position and counts were plotted and fit to a 2-term Gaussian, from which the FWHM was extracted. A precise measure of sperm membrane diameter was obtained by fitting cross-sectional STORM data from ABHD2 to a circle using a least mean squares minimization algorithm(Thomas and Chan, 1989). Hv1 geometry was characterized by binning XZ- or YZ- cross-sectional STORM data into 2D histograms. Peaks from these histograms were used to define the centers of distinct clusters of molecular coordinates; the distance between the centers of mass of these clusters was computed.

**Scanning electron microscopy (SEM).** SEM was performed on fully hydrated sperm cells via graphene protection, as described previously(Wojcik et al., 2015). Briefly, fixed cells on coverglass were stained with 2% uranyl acetate (SPI 02624) in water for 2 hours and thoroughly rinsed. Monolayer CVD graphene on copper foil was obtained from Graphene Supermarket (Calverton, NY). To cover cells with graphene, the hydrated coverslip containing the cells was used to scoop up a graphene-PMMA stack floating on water. The stack was allowed to adhere to the sample for ~10 min in air. To remove PMMA, the sample was dipped in anisole or acetone for 2 min, and rinsed off briefly in isopropyl alcohol. The graphene-covered coverslip was mounted on a standard metallic sample mount with carbon tape, and a small amount of silver colloid paint (Ted Pella 16031) was used to create a conductive bridge between graphene and the sample mount. SEM imaging was performed under standard secondary electron mode on an FEI Quanta 3D FEG system at normal operational vacuum (~$10^{-5}$ torr). Calibration of magnification was verified with a replica of a 2,160 lines/mm waffle-pattern diffraction grating (Ted Pella 604-A).

**Immunogold-labelling and transmission electron microscopy.** After swim-up purification sperm were allowed to sediment. Cells were fixed with PBS/8% PFA for 10 min. After permeabilization with PBS/0.1% saponin for 15 min and blocking with PBS/5% BSA for 45 min, sperm were incubated with a 1:50 dilution of rabbit anti-Hv1 IgG (#AHC001; Alomone Labs, Jerusalem, Israel) in PBS/5% BSA overnight at 4°C. After excessive washing in PBS sperm were incubated for 1 h with goat anti-rabbit antibody conjugated to nanogold (1:40 in PBS/5% BSA; Electron Microscopy Sciences, Hatfield, PA, USA). After three washing steps in PBS, cells were fixed for 15 min with 2.5% glutaraldehyde in PBS. Cells were transferred to agarose and treated for 30 min with 0.5% $OsO_4$ in 0.1 M cacodylate buffer (pH 7.4). After dehydration in a graded ethanol series and propylene oxide the pellet was embedded in Epon 812 resin. After polymerization for 72 h at 65°C, 70 nm thick sections were cut on a Reichert Ultracut-E microtome (Leica, Wetzlar, Germany) and mounted onto formvar-coated nickel grids (Electron Microscopy Sciences, Hatfield, PA, USA). After signal enhancement with the HQ silver enhancement kit (Nanoprobes, Yaphank, NY, USA) for 8 min the grids were treated with 2% uranyl acetate and lead citrate for 4 min, respectively. Samples were examined on a Tecnai 12 transmission electron microscope (FEI, Hillsboro, OR, USA) equipped with an Ultrascan 1000 camera (Gatan, Pleasanton, CA, USA).

**Measurement of the distance between two clusters "A" and "B" in EM cross sections.** The straight distance between the center of each LC and corresponding gold particles in the fibrous sheath (FS) of individual cross sections was measured. Positions of gold particles, which clustered in position "A" or "B" at the 3 plane were averaged to calculate the mean distance of positions "A" (mean A) and "B" (mean B) from each LC. To measure the distance between [mean A] and [mean B], we projected the centers of each LCs to [mean A] and [mean B] from three individual cross sections. Next, three triangles were plotted with defining sides as follows. Triangle 1: **a** = distance between [mean A] and ipsilateral LC, **d** = distance between both LCs (aka flagellar diameter), and **a'** = distance between [mean A] and contralateral LC. The angle between **a** and **d** was defined as **A**. Triangle 2: **b** = distance between [mean B] and ipsilateral LC, **b'** = distance between [mean B] and contralateral LC, and **d**. The angle between **b** and **d** was defined as **B.** Triangle 3: **a**, **b** and **c** = distance between [mean A] and [mean B]. The angle between **a** and **b** was defined as **C.** To determine angles **A** and **B**, the law of cosines was used: $(a')^2= a^2 + d^2 - 2ad\cos(A)$ and $(b')^2= b^2 + d^2 - 2bd\cos(B)$. Since $C= A-B$, we have determined **c** as: $c^2= a^2 + b^2 - 2ab\cos(C)$.

**Simulation of proton efflux via Hv1 and its effect on the intraflagellar pH change.** For the first *in silico* estimation of the global pH rise in the sperm tail (Fig. S7A) we implemented a program in BASIC in which a cylindrical radial diffusion model used a finite-difference-element numerical solution of the diffusion equation, with 5-nm shells and 20-µs time steps. Hydronium diffused at $9.3\times10^{-5}$ $cm^2/s$ (al-Baldawi and Abercrombie, 1992) in the presence of 100 mM MES ($K_D$ = $7.08\times10^{-7}$ M) and a 85 mM of a fixed buffer, with $K_D$ = $9.3\times10^{-7}$ M, as in similar mammalian cells (Swietach et al., 2003). Buffer equilibration was assumed to be instantaneous and buffer diffusion was ignored, because both have no effect on final pH levels reached.

For the second *in silico* estimation of the flagellar pH changes under physiological conditions (Fig. S7B) we used the same program and simulation assuming a starting pH of 6.0, and a temperature of 37 °C rather than 25 °C, which should increase hydronium diffusion by 30% (to $1.212\times10^{-4}$ $cm^2/s$). Assuming that the membrane potential under this condition would be around 0 mV, we modified the appropriate outward current of Fig. S1A (right panel) for human sperm using a temperature coefficient $Q_{10}$ of 2.8 for current amplitudes and $Q_{10}$ of 7 for activation time constants (DeCoursey and Cherny, 1998), corresponding to a current rising to 20 pA with a time constant of 59 ms (Lishko et al., 2010). MES was replaced with 30 mM of a native diffusible buffer with $K_D$ = $2.7\times10^{-8}$ M (Swietach et al., 2003), which would have been replaced in patch clamp experiments with pipette solution.

For the third and final simulation to estimate the steady-state level of pH that might be reached in the neighborhood of a single Hv1 dimer in the presence of buffers (Fig. S7C), we used the point source diffusion equation into a semi-infinite space of equation 6 of (De-la-Rosa et al., 2016), as derived by (Nunogaki and Kasai, 1988) and (Decker and Levitt, 1988):

$$c(r)= c_\infty(1- (a/r)e^{-\lambda(r-a)}), \text{ with } \lambda^2=k_b[B]/D_H$$

Where $c(r)$ is the proton concentration in the presence of a rapid buffer, $c_\infty$ is the bulk proton concentration, $a$ is the capture radius of the channel "sink", $r$ is the radial coordinate indicating the vicinity of a proton channel or a distance from the "sink", $D_H$ is the diffusion coefficient of protons in water, $[B]$ is the concentration of buffer, and $k_b$ is the association rate constant. This equation is highly sensitive to $a$ for hydrogen ions (here assumed to be 2 nm), and depends on buffer concentration $[B]$ and on-rate $k_b$ [$3\times10^{10}$ /mol·s, from (Nunogaki and Kasai, 1988)] as well as the hydronium diffusion rate $D_H$ ($9.3\times10^{-5}$ cm$^2$/s). Note that $c(r)$ achieves diffusional steady state in a few ns ($\tau = r^2/4D_w$ = 2.7 ns at 10 nm), followed by a smaller and slower phase due to buffer equilibration and diffusion (*12*).
